# Polycystin-2 ciliary extracellular vesicle release and targeting

**DOI:** 10.1101/2020.04.21.050690

**Authors:** Juan Wang, Inna A. Nikonorova, Amanda Gu, Paul W. Sternberg, Maureen M. Barr

## Abstract

Extracellular vesicles (EVs) are emerging as a universal means of cell-to-cell communication and hold great potential in diagnostics and regenerative therapies [1]. An urgent need in the field is a fundamental understanding of physiological mechanisms driving EV generation and function. Ciliary EVs act as signaling devices in *Chlamydomonas* and *C. elegans* [2–4]. Mammalian cilia shed EVs to eliminate unwanted receptors [5] or to retract cilia before entering the cell cycle [6]. Here we used our established *C. elegans* model to study sensory-evoked ciliary EV release and targeting using a fluorescently labeled EV cargo polycystin-2 (PKD-2). In *C. elegans* and mammals, the Autosomal Dominant Polycystic Kidney Disease (ADPKD) gene products polycystin-1 and polycystin-2 localize to cilia and EVs, act in the same genetic pathway, and function in a sensory capacity, suggesting ancient conservation [7]. We find that males deposit PKD-2-carrying EVs onto the vulva of the hermaphrodite during mating. We also show that mechanical stimulation triggers release of PKD-2-carrying EVs from cilia. To our knowledge this is the first report of mechanoresponsive nature of the ciliary EV release and of ciliary EV directional transfer from one animal to another animal. Since the polycystins are evolutionarily conserved ciliary EV cargoes, our findings suggest that similar mechanisms for EV release and targeting may occur in other systems and biological contexts.

---

*C. elegans* male mating involves stereotyped behavioral steps including response to hermaphrodite contact, location of the hermaphrodite’s vulva, spicule insertion, and sperm transfer to the hermaphrodite’s uterus [7]. To examine male-hermaphrodite EV-mediated interactions during mating, we paired fluorescently labeled transgenic adult males with unlabeled hermaphrodites for 24 hours (Figure 1A). Male sperm transfer was visualized with MitoTracker dye, whereas ciliary EVs were tracked via the PKD-2::GFP EV cargo protein. In all mated hermaphrodites inseminated with MitoTracker labeled sperm, we observed deposition of male-derived PKD-2::GFP EVs on the hermaphrodite vulvae (Figure 1B-C). No PKD-2::GFP EVs were found inside the hermaphrodite uterus. Location of the male-deposited EVs at the hermaphrodite’s vulva is consistent with the position of a male tail during mating and suggests that EVs were released in the timeframe between successful location of the vulva and retraction of spicules post-copulation. This timeframe represents the closest contact between the male tail and the vulva area of the hermaphrodite, suggesting that the vulva may provide mechanical or chemical cues to stimulate ciliary EV release from the male.

**Figure 1.**
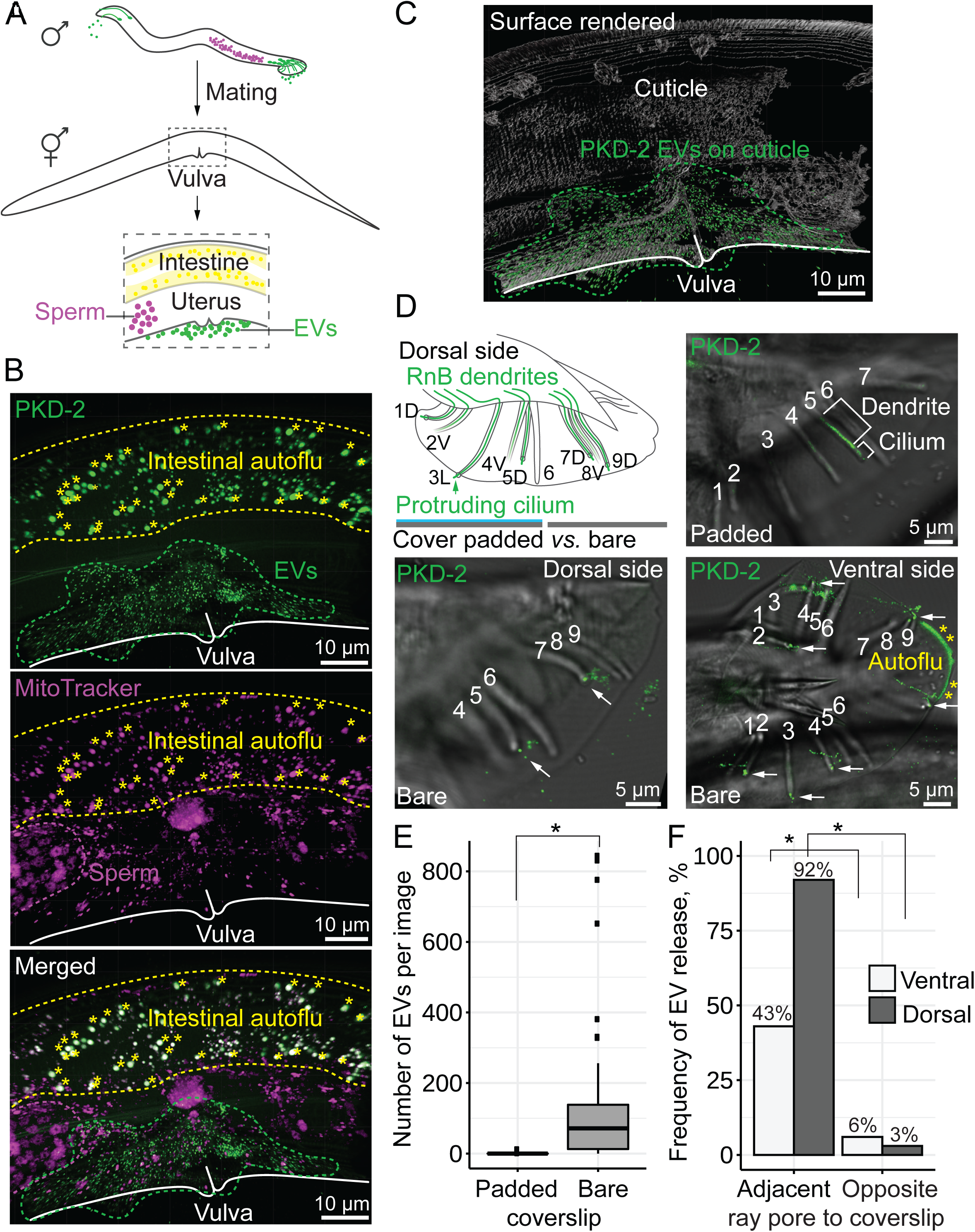
Polycystin-2 extracellular vesicles (EVs) are released upon mechanical stimulation of ciliary tip of male-specific sensory neurons and are directionally transferred to hermaphrodite’s vulva during *C. elegans* mating. (A) Experimental design for tracking the PKD-2::GFP EVs and MitoTracker labeled spermatozoa during mating. Unlabeled hermaphrodites were mated with labeled males and examined under the microscope post-copulation to score for male-derived PKD-2::GFP-labeled EVs and MitoTracker labeled sperm. Insert diagram shows essential anatomical structures of hermaphrodite to aid in interpretation of images on panel B. (B) Projected Z-stack of confocal sections through the inseminated hermaphrodite. The vulva is outlined with solid white; PKD-2 carrying EVs, sperm, and intestine are outlined with green, magenta, and yellow dashed lines, respectively. Autofluorescence from intestinal granules is marked with asterisks; autofluorescence is visible through both channels. (C) Surface rendering of the confocal optical sections from (B) performed by Imaris software, showing that EVs are located on the cuticle around vulva. No PKD-2::GFP EVs were found in the uterus. (D) Diagram of male tail shows identities of sensory ray cilia protruding into environment: 1/5/7/9 protrude from a dorsal side, 2/4/8 protrude from a ventral side of the male tail. Ray 3 is lateral; its cilium protrudes from the edge of the male tail. Ray 6 is a closed ray; its neurons do not express the *pkd-2* gene. Confocal optical sections show EV release (indicated with arrows) upon contact of either dorsal or ventral side of a tail with a bare coverslip. Contact with a padded coverslip rarely triggers EV release. The posterior-most edge of the male fan is autofluorescent (yellow asterisks). (E) Quantitative analysis of ciliary EV release upon contact with either padded or bare coverslip. Each male was first imaged being covered with an agarose-padded coverslip, then the padded coverslip was replaced with a bare coverslip and imaging was repeated for that male. The box plots show median values, top and bottom hinges correspond to the first and third quartiles (the 25th and 75th percentiles), and the whiskers extend to the smallest and the largest value within 1.5 distance of the interquartile range. (F) Quantitative analysis of the propensity of dorsal and ventral cilia to release the PKD-2 EVs when positioned on a side that is either adjacent or opposite to the coverslip. * p-value <0.01, n=39 in the Kruskal-Wallis test for (E); n=56 for dorsal rays, n=42 for ventral rays in the two proportion Z-test for (F).

Living *C. elegans* males release EVs when mounted between an agarose-layered slide and a bare glass coverslip [3], with EVs usually floating close to the surface of the coverslip. To test the hypothesis that the coverslip might mechanically stimulate the EV release, we padded the coverslip with a thin layer of agarose gel to reduce mechanical stimulation (see Supplemental methods for details). The use of the agarose padded coverslip drastically reduced number of released EVs (Figure 1D-E), indicating that the soft surface did not stimulate PKD-2::GFP EV release. We then proceeded to replace the padded coverslip with a new bare coverslip and observed abundant release of the ciliary EVs (Figure 1D-E). These data suggest that mechanical stimulation from the bare coverslip triggers the PKD-2::GFP EV release.

Anatomically, the male tail has ventrally and dorsally positioned rays [8]. Each ray contains a B-type cilium that releases EVs and protrudes through a cuticular pore into the environment (Figure 1D). Our analysis revealed that EVs were more frequently released from ray cilia tips that were adjacent to the bare coverslip (Figure 1D, F). PKD-2::GFP EVs were also released from the hook B-type neuron (HOB) when ventral side of a male tail faced the bare coverslip (Figure S1). Statistical analysis of frequencies of the EV release events from dorsal and ventral sides indicated that the EV release corresponded with the position of the ray pore and cilium against the bare coverslip (Figure 1F and Table S1). This data support the hypothesis that mechanical stimulation triggers PKD-2::GFP EV release. We propose that this mechanoresponsive nature of PKD-2 EV release *in situ* may act during male-hermaphrodite mating *in vivo*.

Our finding that the male directly deposited PKD-2::GFP-labeled EVs to the hermaphrodite’s vulva suggests that the EV release is an activity-evoked event rather than a constitutive process. This dynamic implies that the PKD-2 protein should be actively and rapidly transported toward ciliary tip during appropriate mechanical stimulation, as suggested by our data. An alternative interpretation is that, in addition to mechanical stimulation, EV release might be triggered by the chemical landscape of the bare glass coverslip. We consider the latter scenario unlikely because chemical cues from the glass surface would likely be overridden by the ion-rich phosphate-buffered solution in which worms were mounted during imaging sessions. We predict that during mating, both mechanical and chemical cues might work in concert to trigger and target male-derived ciliary EVs specifically to the hermaphrodite’s vulva.

Our previous studies show that isolated PKD-2-carrying EVs elicit male tail-chasing behavior and function in animal-to-animal communication [3, 9]. Here we discovered that male-derived EVs are targeted to the vulva of its mating partner *in vivo*. However, EV cargo content and the exact function of this targeting cannot be inferred from these studies. Transcriptional profiling of these EV-releasing neurons revealed a large variety of adhesive membrane proteins, cellular stress components, and innate immunity modulators, including antimicrobial peptide EV cargo [10]. Since EVs are reported to carry a variety of signaling molecules and enzymes, possibilities for their function are numerous. EVs might prevent males from mating with inseminated hermaphrodites, protect inseminated hermaphrodites from microbial invasion, or aid in the mating process by precisely marking the vulva site. Finally, EV release may maintain cilium structure and function, akin to photoreceptor disc shedding in the vertebrate eye. In conclusion, this study opens a new venue for exploring ciliary EV function in inter-organismal communication and in reproductive biology.

## SUPPLEMENTAL INFORMATION

Supplemental information includes one table, one supplemental figure, and experimental procedures.

## AUTHOR CONTRIBUTIONS

Study conception and design: J.W., P.W.S., M.M.B. Acquisition of data: J.W. Analysis and interpretation of data: J.W., A.G., I.A.N., M.M.B. Writing of manuscript: J.W., I.A.N., M.M.B.

## ACKNOWLEDGEMENTS

This work was supported by grants from Kansas PKD Research and Translation Core Center, P30 DK 106912 to J.W. and National Institutes of Health (NIH) awards DK059418 and DK116606 to M.M.B. We thank Noriko Kane-Goldsmith for assistance with confocal microscopy; Gloria Androwski and Helen Ushakov for excellent technical assistance; Barr labmates and the Rutgers *C. elegans* community for feedback and constructive criticism throughout this project; Joel Rosenbaum, Robert O’Hagan, Natalia Morsci, and the three anonymous reviewers for insightful suggestions on this manuscript; and WormBase. We also thank the *Caenorhabditis* Genetics Center (CGC) for strains. The CGC is supported by the National Institutes of Health - Office of Research Infrastructure Programs (P40OD010440).

## Supplemental Information

**Supplemental Figure S1.**
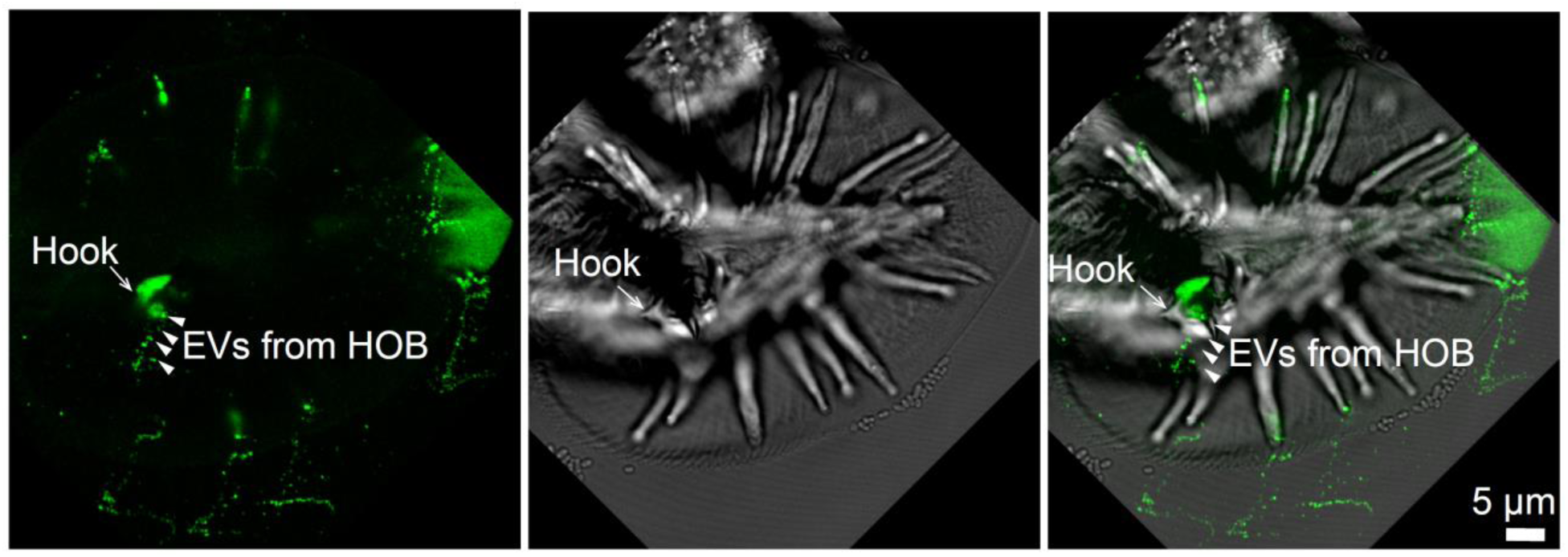
Release of the PKD-2::GFP-labeled EVs from the HOB sensory cilium of the male tail. Optical sections through *C. elegans* male tail positioned ventrally to the bare coverslip. The hook structure with HOB sensory cilium is labeled with arrow, PKD-2::GFP EVs released from the HOB cilium are labeled with arrowheads.

**Supplemental Table S1.**
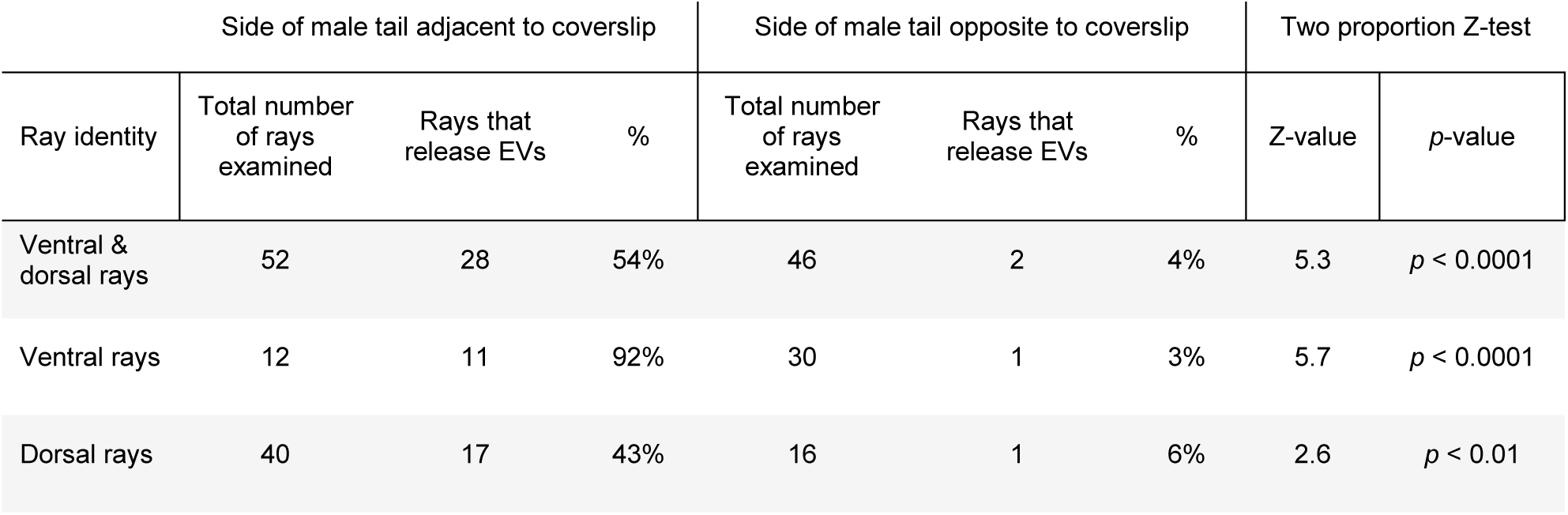
Statistical analysis of the frequencies of PKD-2::GFP EV releasing events from ray B type ciliated neurons of the male tail when positioned either adjacent or opposite to the coverslip.

| Table S1. Side of male tail adjacent to coverslip |  |  |  | Side of male tail opposite to coverslip |  |  | Two proportion Z-test |  |
| --- | --- | --- | --- | --- | --- | --- | --- | --- |
| Ray identity | Total number of rays examined | Rays that release EVs | % | Total number of rays examined | Rays that release EVs | % | Z-value | *p*-value |
| Ventral & dorsal rays | 52 | 28 | 54% | 46 | 2 | 4% | 5.3 | *p* < 0.0001 |
| Ventral rays | 12 | 11 | 92% | 30 | 1 | 3% | 5.7 | *p* < 0.0001 |
| Dorsal rays | 40 | 17 | 43% | 16 | 1 | 6% | 2.6 | *p* < 0.01 |

## Supplemental Experimental Procedures

### Strains

All *C. elegans* strains were maintained under standard conditions as described in [S1]. The mating assay used uncoordinated hermaphrodites of the CB369 (*unc-51* (e369)) strain to serve as easy mating targets for males. The males were of the PT443 strain (*myIs1*[P*pkd-2*::PKD2::GFP+P*unc-122*::GFP] I; *pkd-2(sy606)* IV; *him-5(e1490)* V) that carries a loss-of-function *pkd-2* allele together with an integrated rescuing array encoding GFP-labeled PKD-2 protein [S2].

Scoring of the PKD-2 EV release was performed using the PT3112 strain (*pha-1(e2123)* III; *him-5(e1490)* V, *myIs4*[P*pkd-2*::PKD2::GFP+P*unc-122*::GFP] V; *myEx888*[CIL-7::tagRFP + pBX1])[This work].

### Mating assay tracking sperm and EV transfer

Male worms were pre-stained by soaking in 10 μM MitoTracker Ted CMXRos dye (ThermoFisher Scientific #M7512) buffered with the isosmotic M9 buffer (22 mM KH_2_PO_4_, 43 mM Na_2_HPO_4_, 85 mM NaCl, 1 mM MgSO_4_) for 7 hours in dark to label their spermatozoa [S3]. Prior to introduction to the hermaphrodites, males were allowed to crawl on a clean plate to remove any excess of the dye. Mating was conducted on an agar plate with a bacterial lawn of 1 cm in diameter generated with 15 μl of overnight OP50 *E. coli* culture. Ten unstained L4 *unc-51 (e369)* hermaphrodites were placed together with fifty pre-stained males and allowed to mate for 24 hours in the dark at 22°C. Following mating, hermaphrodites were mounted in a 0.5 μl drop of 10 mM levamisole (prepared in the M9 buffer) placed on a 5% agarose pad (prepared with ultra-pure water and agarose, Sigma #A9539) for further imaging.

### Mounting of males for quantitative scoring of the EV release

For regular imaging of the EV release, males were mounted on agarose layered glass slides in the same way as described above for the hermaphrodites [S4]. In order to diminish mechanical stimulation and test the hypothesis about mechanoresponsive EV release, we developed the double agarose sandwich protocol, where not only the microscope glass slide is layered with 5% agarose, but the coverslip also has a thin padding made from agarose gel. To prepare the padded coverslips with a thinnest possible agarose pad, 20 µl of 5% agarose gel were dropped on a pre-heated to 90°C coverslip and pressed with another coverslip. Then, the coverslip-agarose sandwich was cooled to ambient temperature and was carefully separated so that only one piece of the coverslip left covered with the agarose gel padding. Imaging of male worms mounted with the agarose-padded coverslips was performed in 1 mM levamisole. After each imaging session, the agarose padded coverslip was replaced with a bare coverslip to conduct a second imaging session on the same male to score its response to a bare coverslip. Both imaging sessions were performed within a 30-minute timeframe.

### Fluorescent imaging and quantification

Images were acquired using Zeiss LSM880 confocal microscope with Airyscan high-resolution detector. Image processing included Airyscan processing performed with the accompanying Zeiss software ZEN 2 (Blue version). Surface rendering of the hermaphrodite images was obtained with Imaris software. PKD-2::GFP EVs were quantified using ZEN Blue imaging analysis software.

### Statistical analysis

Non-parametric Kruskal-Wallis test was used to test the hypothesis that the double agarose sandwich mounting protocol results in significantly different number of EVs released from cilia of a male tail.

Two-proportion Z-test was used to test the hypothesis that frequencies of EV release events from male ray cilia facing or not facing the bare coverslip are significantly different (Table 1). Lateral ray 3 was also producing EVs but was excluded from the analysis as its position relative to coverslip could not be established unequivocally.

